# Genomic differentiation within East Asian *Helicobacter pylori*

**DOI:** 10.1101/2021.06.05.447026

**Authors:** Yuanhai You, Kaisa Thorell, Lihua He, Koji Yahara, Yoshio Yamaoka, Jeong-Heon Cha, Kazunari Murakami, Yukako Katsura, TEAMHp, Ichizo Kobayashi, Daniel Falush, Jianzhong Zhang

## Abstract

The East Asian region, including China, Japan and Korea, accounts for half of gastric cancer deaths. However, different areas have contrasting gastric cancer incidence and the population structure of *Helicobacter pylori* in this ethnically diverse region is yet unknown. We aimed to investigate genomic differences in *H. pylori* between these areas to identify sequence polymorphisms associated with increased cancer risk.

We analysed 381 *H. pylori* genomes collected from different areas of the three countries using phylogenetic and population genetic tools to characterize population differentiation. The functional consequences of Single Nucleotide Polymorphisms (SNPs) with a highest fixation index (Fst) between subpopulations were examined by mapping amino-acid changes on 3D protein structure, solved or modelled.

329/381 genomes belonged to the previously identified hspEAsia population indicating that import of bacteria from other regions of the world has been uncommon. Seven sub-regional clusters were found within hspEAsia, related to sub-populations with various ethnicities, geographies and gastric cancer risks. Sub-population-specific amino-acid changes were found in multi-drug exporters (*hefC*), transporters (*frpB-4*), outer membrane proteins (*hopI*), and several genes involved in host interaction, such as catalase, involved in H_2_O_2_ entrance, and a flagellin site mimicking host glycosylation. Several of the top hits including *frpB-4, hefC, alpB*/*hopB*, and *hofC*. were also differentiated within the Americas, indicating that a handful of genes may be key to local geographic adaptation.

*H. pylori* within East Asia are not homogeneous but have become differentiated geographically at multiple loci that have facilitated adaptation to local conditions and hosts. This has important implications for further evaluation of these changes in relation to the varying gastric cancer incidence between geographical areas in this region.

## Introduction

*Helicobacter pylori* has co-evolved with human beings for at least 100,000 years and has strong association with the occurrence of gastric (stomach) cancer ^1^. *H. pylori* are transmitted most effectively within households and have therefore experienced low migration rates compared to many other members of the human microbial flora. As a result, we can expect the pattern of differentiation to reflect historical human migration patterns. The population structure of *H. pylori* worldwide has been classified into seven major groups that indeed correlate with ancient human migrations ^2^, of which the hpEastAsia includes at least three subgroups: hspEAsia, hspAmerind and hspMaori. The hspEAsia subgroup is thought to be ubiquitous within East Asian countries with high gastric cancer incidence, including China, Japan and Korea. These countries together make up 1/5 of the world population but account for half the global mortality from the disease ^3, 4^. The hspEAsia strains have documented higher virulence than other subpopulations and have diverged from the Western strains in several proteins including virulence factors ^5, 6^. Many genes have also diverged *within* this region ^5^.

The prevalence of gastric cancer show geographic and ethnic variations also within East Asia. Some north and southeast areas of China such as Fujian have higher incidence ^7^, whereas some west regions, such as Yunnan, have low incidence (**Figure 1**) ^8, 9^. China also shows diverse distribution of population ethnicities; in southwest Yunnan province more than twenty ethnicities exist. South Korea has been reported with the highest incidence in the world ^10, 11^ and similar is true for the Japanese main islands (including Hokkaido) ^12^, while in Okinawa, where the ethnic composition is different, the incidence is low ^13^. Previous studies on the high-risk regions have suggested that diet, lifestyle and *H. pylori* properties may contribute to the high risk. However, despite the complex human migrations and evolutionary history in these areas, variation of *H. pylori* within the hspEAsia subpopulation has been poorly explored.

**Figure.1.**
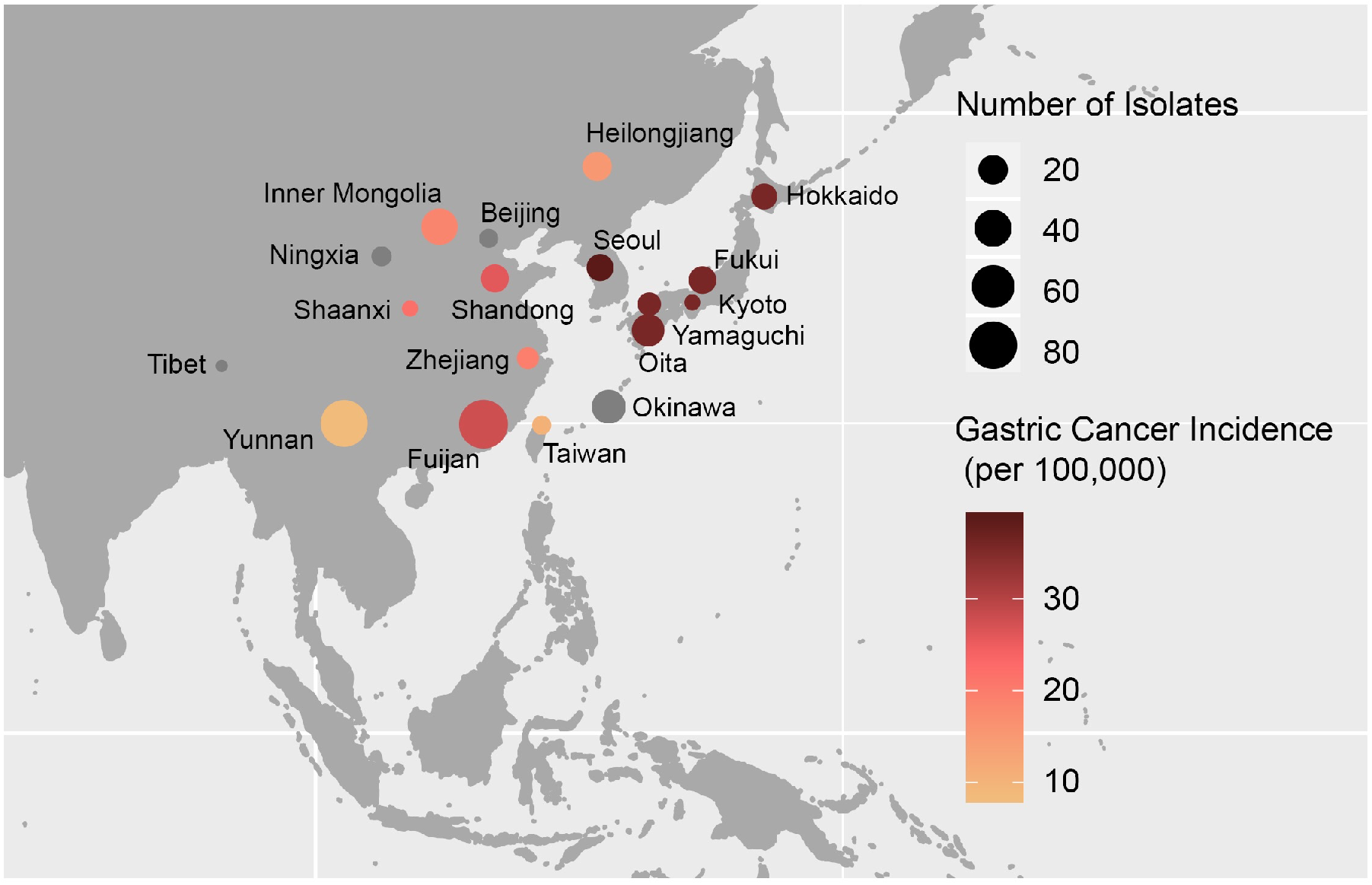
Geographic distribution of the East Asian *H. pylori* strain dataset of this study. Shade of the circle fill indicates gastric cancer incidence. Grey fill indicates missing incidence data.

In the present work, we analysed a collection of genomes of *H. pylori* strains from various places in these three countries. Our phylogenetic and population genetic analysis revealed presence of pronounced regional and ethnical population structure within hspEAsia, and specific sequence differences differentiating these subpopulations. Most of these highly region-specific variants were found on proteins involved in host interaction. Furthermore, placement of the variants on protein structure provided insight into molecular mechanisms underlying regional adaptation.

## Methods

### Strain collection across East Asia

We collected 357 *H. pylori* isolates from China, Japan and South Korea, isolated between 1999 and 2018, including 11 provinces of China and 6 regions of Japan (**Supplementary Tables S1-S2**, **Figure 1**). We use data from 2014 to show regional risk levels in China and compared it with data from Japan and Korea ^7, 8, 10–13^. Our sample of 77 Yunnan isolates includes strains from 4 ethnic minorities and the Han majority individuals.

### Genome sequencing

Genomic DNA was extracted using the Qiagen DNeasy Mini Kit and genomes were sequenced using Illumina or PacBio sequencers. Quality assessment of genomic DNA, SMRTbell library preparation, and data evaluation were performed for PacBio RS II sequencing. Sequences were deposited into GenBank with BioProject number PRJNA482300. Combining publicly available genomes with these newly sequenced genomes, we collected a dataset consisting of 381 *H. pylori* genomes for further analysis (**Supplementary Table S2**).

### Genomic comparison

We used Snippy ^14^ to perform a whole genome alignment with XZ274 as reference genome and extracted 225 942 variable core genome sites. For the phylogenetic analysis, the dataset of 381 strains were combined with 1-3 reference sequences for each of the major *H. pylori* populations (**Supplementary Table 3**).

### Population structure and phylogenetic analysis

The concatenated whole genome SNPs dataset was also used to prepare the haplotype data for ChromoPainter and fineSTRUCTURE analyses ^15, 16^. 13 highly clonal sequences were removed prior to the analysis, resulting in a comparison of 368 genomes. The recombination rate was set to 0.000001 per site/base according to previous studies ^16^. Using each genome as both donor and recipient haplotypes, we used ChromoPainter to calculate the number of genetic chunks exported from a donor to a recipient and generated a co-ancestry matrix. Then fineSTRUCTURE was run by setting the burn-in and Markov chain Monte Carlo (MCMC) chain of 100,000 iterations to generate clusters for all the individual strains. We also generated a phylogenetic tree using FastTree ^17^, which was labelled using iTol ^18^.

### Definition of subgroups and calculation of Fst between subgroups

Subgroups were defined according to the population structure analysed by fineSTRUCTURE. We named the subgroups assigned by fineSTRUCTURE as “Sg” (**Supplementary Table S2**). To identify the SNPs attributed to the divergence of subgroups more accurately, we used a more stringent definition of the subgroups. Only those isolates assigned into a singular cluster in the fineSTRUCTURE tree were defined to form a subgroup. For example, for China southwest isolates, only those from Yunnan Mosuo and Pumi ethnicities, which clustered into a singular clade, were defined as a subgroup, “YunnanMP”. For China southeast, only those from Fujian Changle that clustered into a singular clade were defined as a subgroup, “Fuijan”.

For each SNP site, we calculated a fixation index (Fst) for each subgroup such that Fst (sg1) = pairwise calculation of Fst of sg1 versus all other subgroups. We also compared isolates from China with those from Japan and South Korea and the two lower-incidence regions, Yunnan and Okinawa, versus high incidence regions (the remainder of hspEAsia strains).

### Mapping high-Fst SNPs on protein structure

We located each of the SNPs with the highest Fst values on the reference genome of XZ274, a Tibetan strain, to identify its gene and effect on amino-acid sequence. If the gene was missing or atypical in this strain, we used strain F57, a Japanese strain, or 26695, a European strain, instead as shown in Table S4. We mapped the amino acids on solved *H. pylori* protein structure or on protein structure homology-modelled by SwissModel and its repository for 26695 (https://swissmodel.expasy.org/repository?start=0&rows=50&query=Helicobacter+pylori+26695). We analysed and presented them by PyMOL ^19^.

## Results

### *East Asian* H. pylori *population structure is associated with geography and host ethnicity*

To analyse the genetic structure of *H. pylori* in China, Japan and South Korea, we constructed a phylogenetic tree (**Figure 2A**) and clustered the strains using fineSTRUCTURE^15, 16^ (**Figure 2B**), and constructed a phylogenetic tree (**Figure 2A**). The two methods gave broadly concordant clustering and indicated differentiation at multiple scales. By including reference genomes from the other main *H. pylori* populations, we found that the 52 most differentiated strains do not belong to hspEAsia and have had distinct evolutionary histories. One main cluster of these comprised isolates from individuals of Mongolian ethnicity (Sg4, **Figure 2B**), which in the tree grouped between the references of hspIndigenousAmerica (previously called hspAmerind) and hpAsia2 populations. We also identified a cluster of Okinawan strains diverging after hpEurope, and before hpAsia2 and hpEastAsia (**Figure 2A**), also shown in the lowest line of the co-ancestry matrix (**Figure 2B**). This likely corresponds to Group C in MLST and STRUCTURE analysis of Okinawan strains ^20^. Because these populations are best analyzed in the context of broader regional variation, we focused our remaining analysis on variation within the 329 hspEAsia isolates in our sample.

**Figure 2.**
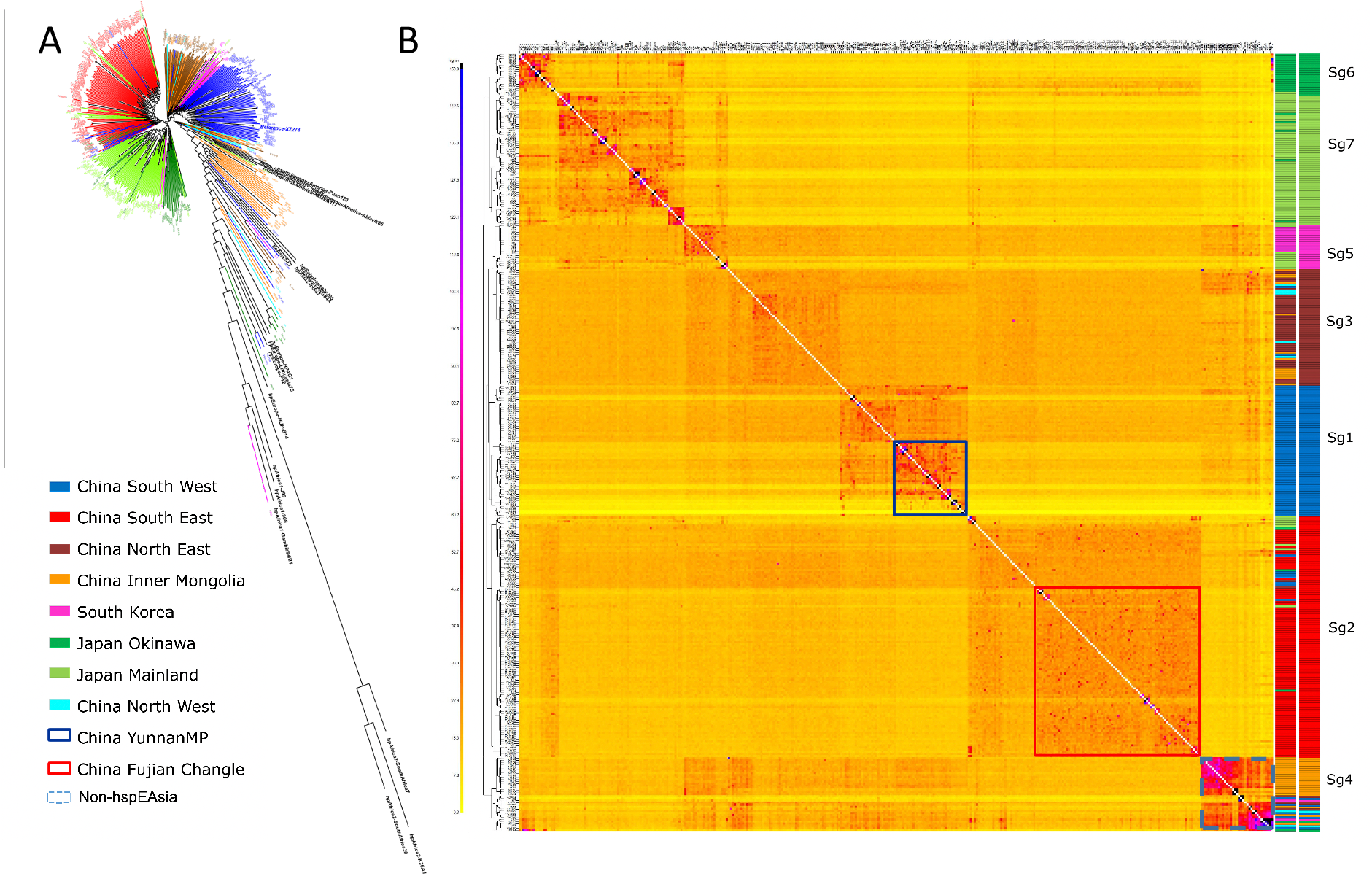
Phylogenetic and population genetic analysis of East Asia *H. pylori* strains. A) Phylogenetic tree of East Asia genomes combined with reference sequences from global *H. pylori* populations. B) Co-ancestry analysis of East Asia isolates after removing highly clonal isolates

HspEAsia isolates are relatively homogeneous but show fine-scale differentiation that is strongly correlated with geography. The majority of Japanese isolates clustered together (Sg7), with Okinawan hspEAsia strains forming a distinct subpopulation (Sg6). The latter likely corresponds to Group A in MLST ^20^. Two strains from Hokkaido cluster with the Sg6 Okinawan strains. This is likely related to the evolutionary structure of Japanese people and of the Okinawa people ^21^. The native ethnicity remains in Hokkaido (as Ainu ethnicity) and in Okinawa.

South Korean isolates also form a distinct subpopulation, which cluster together with isolates from the Northeast of China, concordant with its geographic location. A few Japanese strains are in the Korean cluster while a few Korean strains are in the main Japanese cluster and branching around the same time as the Okinawa cluster (**Figure 2AB**). This may reflect immigration from Korea to Northern Kyushu and Okinawa. Within China, there is clear differentiation between the Northeast, Southeast and Southwest areas. Within the Southwest, a subpopulation containing strains from Yunnan Mosuo and Pumi ethnicities could also be observed. A Tibetan isolate clustered with the Pumi population, consistent with their migration and mixture history. Within the Southeast, there was also a sub-population (Sg2), specific to Fujian Changle, a region of high gastric cancer incidence (**Figure 2, Supplementary table S1**). The Southeast cluster also includes multiple clusters of Japanese/Okinawa strains (**Figure 2AB**) likely reflecting the immigration to Japan/Okinawa especially frequent since the advent of rice-paddy cultivation.

### Many genetic variants show strong differentiation by subpopulations

In order to explore the genetic basis of local differentiation, we calculated fixation index, Fst, between the hspEAsia subpopulations identified by fineSTRUCTURE. To obtain more reliable SNPs associated with subgroup separation, we used the YunnanMP sub-cluster of Sg1 and Fujian sub-cluster of Sg2, along with the remaining 5 subgroups defined by the fineSTRUCTURE analysis. For most of the subpopulations, more than 99% of SNPs were weakly differentiated, with Fst less than 0.3 (**Supplementary Figure S1**). The Korean subpopulation had a smaller sample size and showed the weakest Fst values. Both single nucleotide polymorphisms (SNPs) and clustered nucleotide polymorphisms (CNPs), likely resulting from recombination between diverged sequence groups, were found.

To functionally interpret differentiation between the populations, we focused on the SNPs with the top 20 Fst values in each subgroup and then removed SNPs with Fst < 0.5. This resulted in a list of 56 genes, several of which occurred more than once (**Supplementary Table 4, bold**). In the main text, we focus on the genes with the strongest evidence for differentiation. Six of these had one SNP with Fst>0.6 and appeared in at least two top 20 lists. Another 6 had at least 2 SNPs with Fst>0.6. Of these HofC has multiple non-synonymous SNPs with Fst up to 0.86, marking it out as also being a particularly strong candidate for being differentiated by natural selection.

To predict the effect of differentiated variants on *H. pylori* biology and pathogenesis, we functionally annotated the genes and interpreted the impact of differentiated amino acids on the protein structures, solved or based on homology modelling. The majority of genes containing the most differentiated SNPs could be grouped into four major categories; (i) transporters, (ii) outer membrane proteins, (iii) metabolism and (iv) host interaction.

#### (i) Transporters

##### The hef multidrug efflux pump

*H. pylori* carries 4 gene clusters that each encodes for a set of RND superfamily of multidrug efflux pump corresponding to TolC-AcrA-AcrB of *E. coli* (**Figure 3**). They pump out endogenous bile salts and ceragenins as well as various antibiotics ^22–24^. The single most differentiated SNP in our analysis is in *hefC* (HP0607) in the YunnanMP subpopulation, with an Fst=0.89. This N86S is also the most differentiated between the lower cancer incidence regions (Yunnan and Okinawa) and the remainder. This residue corresponds to the gate to channel III for planar aromatic cations in *E. coli* homolog ^25^ and the regional adaptation may therefore remodel this gate to export chemicals of this type at different ratios. Also, the outer component of the efflux pump, HefD, showed YunnanMP specific residues in the equatorial domain that may be involved in the interaction with the inner component to open the aperture.

**Figure 3.**
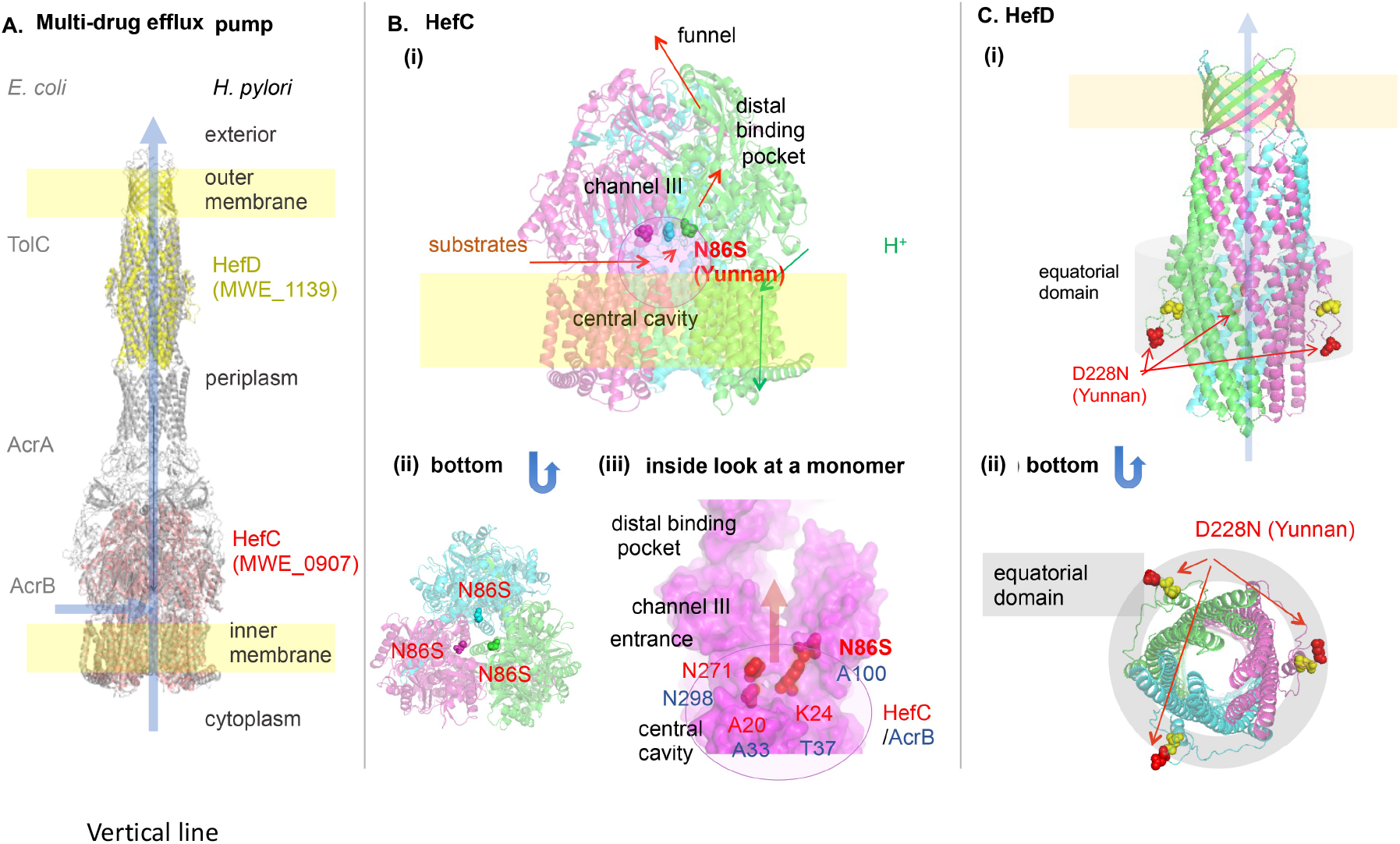
Inferred structure and differentiated amino acid sites of Multidrug efflux pump related proteins. A) TolC-AcrA-AcrB of *E. coli* and its *H. pylori* homologs. B) HefC (MWE_0907), modelled on PDB 3W9I (MexB of *Pseudomonas aeruginosa*). (iii) HefC modelled on PDB 3AOD (AcrB of *E. coli*). Yunnan-specific N86S is at the entrance of channel III. C) HefD (MWE_1139), modelled on PDB 5BUN (ST50 from *Salmonella enterica* subsp. enterica serovar Typhi). Yellow spheres indicate Mongol-differentiated E219D.

##### The TonB-dependent nickel importer frpB-4

A gene containing multiple highly differentiated SNPs, especially in North Eastern China, is *frpB-4* (HP1512), encoding for an outer membrane transporter of nickel of some form ^26^. It is a member of TonB-dependent transporter family, which forms a trimer of 22-stranded beta barrels each filled with a ‘plug’ (**Figure 4A,B**). *H. pylori* reference strain 26695 carries four *frpB* homologs: *frpB-1* (HP0876), *frpB-2/3* (HP0916/5), and *frpB-4* (HP1512).

**Figure 4.**
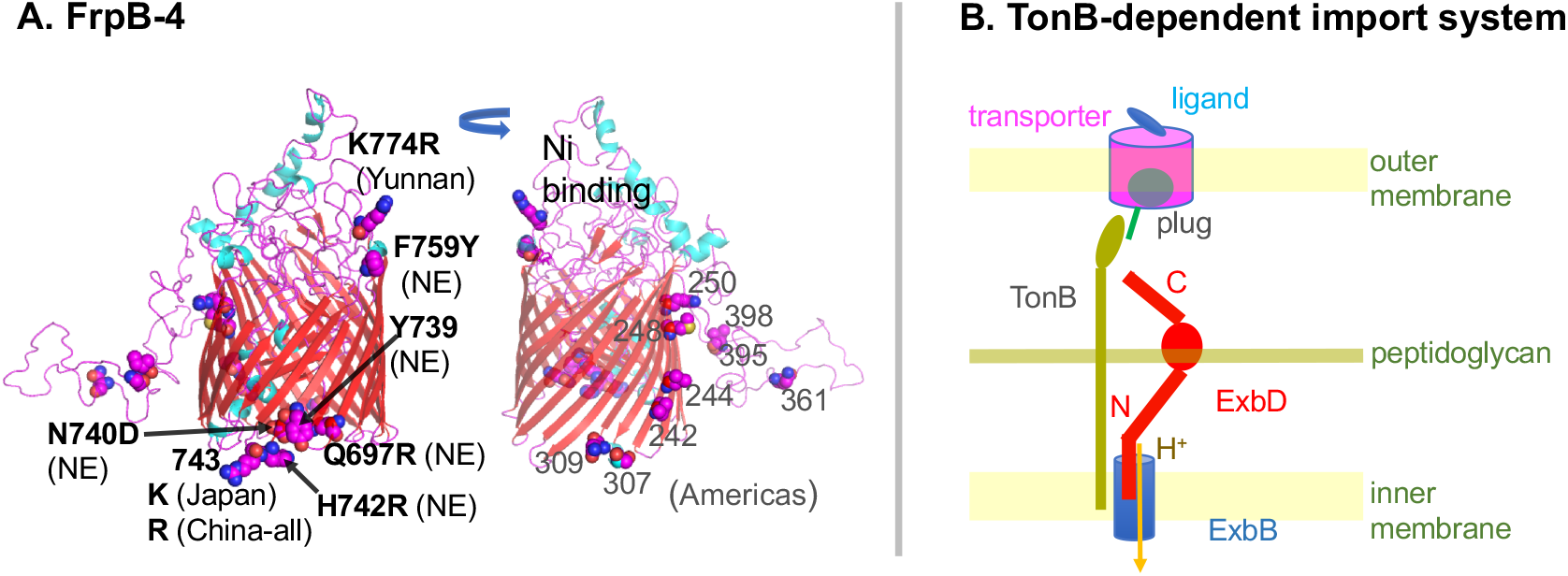
(A) FrpB-4 (MWE_17OO) modelled on PDB 4AIQ (FrpB of *N. meningitidis*) including Americas-specific sites. (B) TonB-dependent transport system. When the transporter in the outer membrane catches a ligand, it pushes out its plug for TonB to pull and open the channel. This movement of TonB is energized by ExbBD motor in the inner membrane driven by H^+^ flow.

Ligand binding lets TonB change the conformation of the plug, which opens a channel, a process energized by the ExbBD proton-driven motor ^27^ (**Figure 4B**). Four China North East-specific Fst sites cluster (Fig. 4D), with three sites (739, 740, 742) presumably representing a CNP (clustered nucleotide polymorphisms). The next residue (743) distinguishes between Japan (K) and China (J). Northeast China-differentiated F759Y (F for the remainder and Y for Northeast China) and YunnanMP-differentiated K774R are situated above the barrel in the model and may interact with the ligand. Various regions of the Americas show region-specific amino-acid changes in other areas of the protein ^28^, out of which three are predicted to be in the decoy loop. Taken together, these changes may affect nickel transport and, consequently, urease activity, since the urease enzyme requires nickel for acid acclimation ^29^. The changes could be related to regional differences in host nickel metabolism and stomach acidity.

##### Other transporters

Hof proteins (*Helicobacter-specific* outer membrane protein family ^30^) are 18-stranded β-barrels homologous to Occ family of *Pseudomonas* and *Campylobacter jejuni* MOMP (major outer membrane protein) involved in passive diffusion of cations including antibiotics and in adhesion ^31–33^. HofC (HP0486), required for *H. pylori* colonization in mice ^34^, contains the most differentiated SNPs in Fujian, a region of high cancer incidence with Fst=0.86. The gene is highly variable in global strains and shows many America-differentiated SNPs and region-differentiated SNPs within the Americas ^29^, within one narrow region. Fujian-differentiated D186S in HP0486 (166 in MWE_0556) lies at a distance from them. Fujian-differentiated V9A is in its signal peptide.

OppD (HP0250), the cytoplasmic subunit of the oligopeptide ABC transporter, has YunnanMP-differentiated KG306R within the ATP binding Walker motif A (**Figure 5A**).

**Figure 5.**
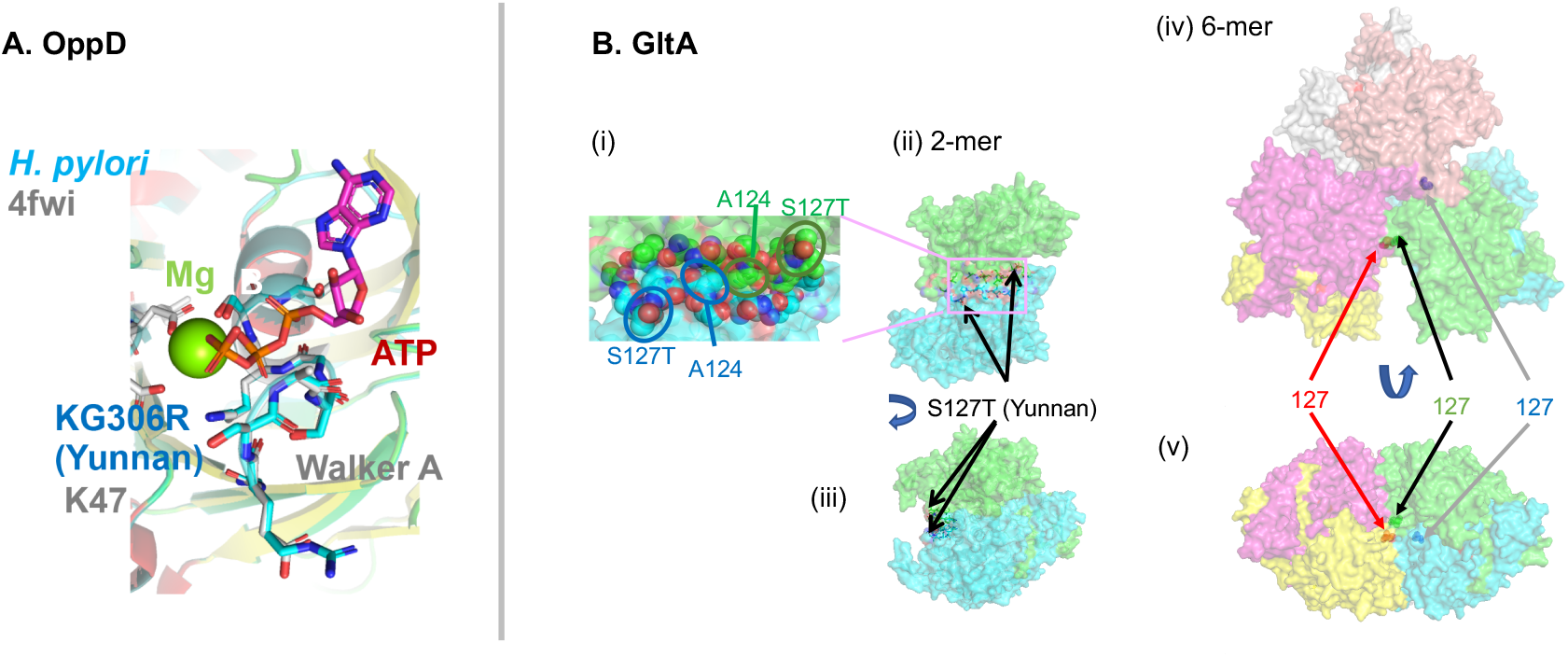
A) OppD (MWE_0327), ATP binding subunit of an oligopeptide ABC transporter as modelled on PDB 4FWI (*Thermoanaerobacter tengcongensis*). B) Citrate synthase, GltA (i) (ii) (iii) MWE_1570 modelled on PDB 3MSU (*Francisella tularensis* homolog). (iv) (v) On PDB 2h12 (*Acetobacter aceti*).

#### (ii) Outer membrane proteins

HopB/ AlpB/ Omp21 (HP0913), an adhesin of the Hop family required for colonization, carries Fujian-differentiated N289D and N286H. According to a previous study, 24 polymorphic sites within 49 bp in AlpB are enriched for Asian ancestry in hspEuropeColombia and 32 polymorphic sites within 65 bp were enriched for Asian ancestry in hspAfrica1Nicaragua populations ^29^.

Another member of the Hop family of outer membrane proteins, HopI (HP1156), has a site (467) that distinguishes between Japan (H) and China-all (D) and YunnanMP-differentiated V633L.

#### (iii) Central Metabolism

The region-differentiated amino-acid changes involve a handful of key metabolic enzymes. Citrate synthase GltA (HP0026, MWE_1570), is the first enzyme in the TCA cycle catalysing the conversion of acetyl-CoA and oxaloacetate to citrate. Yunnan-differentiated S127T is located between the two identical monomers and is likely involved in their association as well as in dimer-dimer association to form a 6-mer (**Figure 5B**). The differentiated SNP might change the quaternary structure. Mutation of A124 in this interface was found in experimental evolution in *E. coli*^35^. In addition to GltA, two other key metabolic enzymes, PorB (HP1111), a subunit of pyruvate:ferredoxin oxidoreductase, and FixP (HP0147, MWE_0216), a subunit of cytochrome c oxidase had high Fst values in YunnanMP. PorB, a key enzyme in the microaerophilic metabolism of *H. pylori*, converts pyruvate to acetyl-CoA, the substrate of GltA, and FixP is a component in aerobic respiration. The Yunnan-differentiated residue is at the proton entrance (Supplementary Figure 4C). Together, these changes might affect the metabolic capacity of this regional *H. pylori* subpopulation.

#### (iv) Host interaction

In addition to the genes listed above, region-specific non-synonymous variants are present in several genes that are annotated as known virulence or host interaction factors.

Catalase KatA (HP0875) (**Figure 6A)** detoxifies H2O2 generated by host immune cells. It also binds host vitronectin, thereby protecting against complement-mediated killing ^36^. The preferred route for H_2_O_2_ is the channel S451-D109-H56-heme (**Fig. 6A (ii))** ^37^. YunnanMP-differentiated P160H by the entrance S451 drastically changes local conformation and surface electric charge. Fujian-differentiated N248D with −1 change in the electric charge takes place near the dimer- dimer interface (**Figure 6A (i)**) likely changing their interaction.

**Figure 6.**
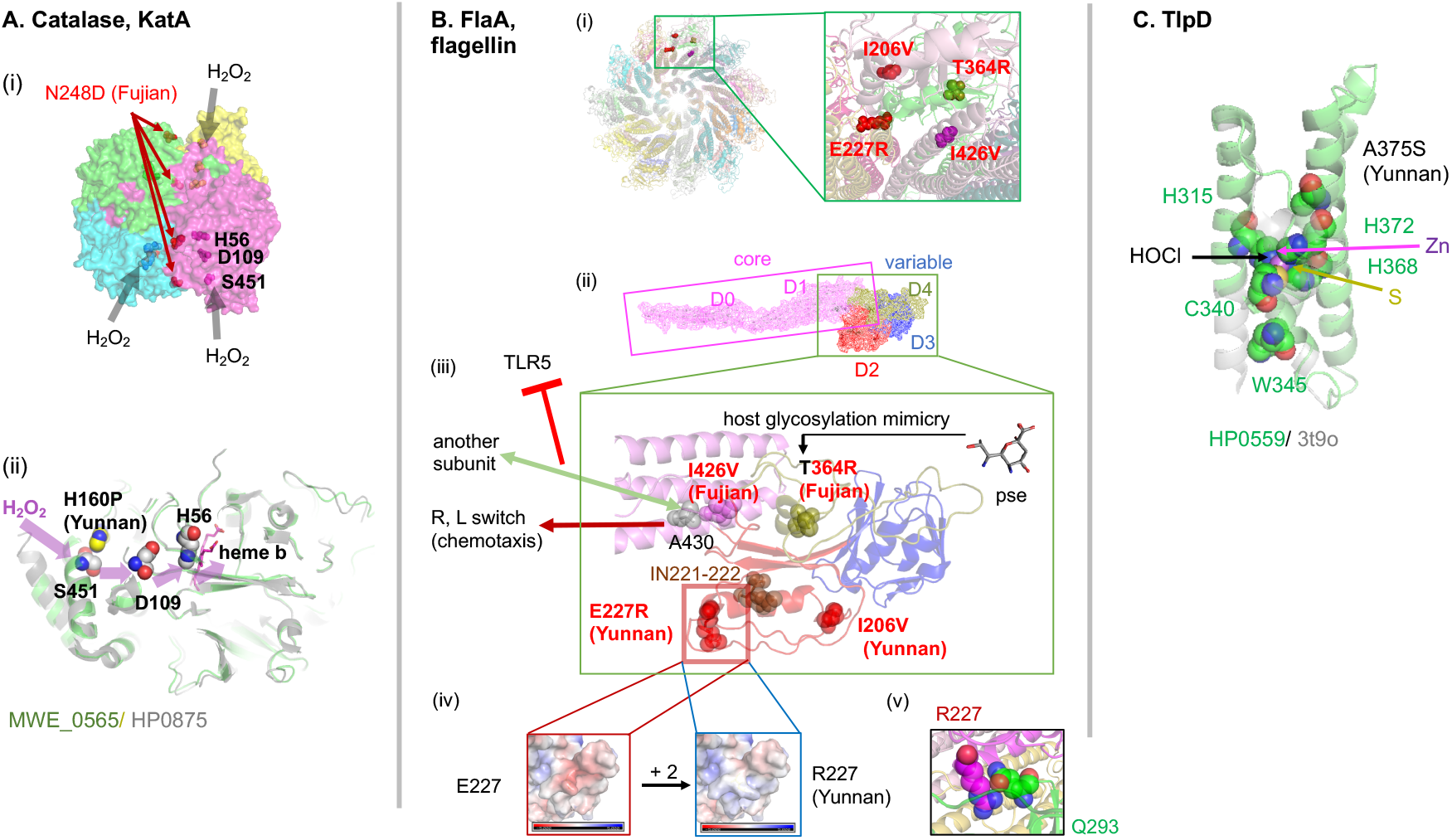
A) Catalase KatA (HP0875, PDB 2A9E). (i) Dimer of dimer. (ii) Channel for H_2_O_2_. Yunnan-specific P160H by its entrance S451 drastically changes local conformation and surface electric charge (mutagenesis in PyMOL). B) Flagellin, FlaA. (i) A view from the distal end of the 22-mer model of FlaA (MWE_0913) on G508A mutant of *Campylobacter jejuni* homolog (PDB 6×80) with 4 population-specific amino-acid changes in two interacting monomers. (ii) Monomer with 5 domains. (iii) Population-specific amino-acid changes. pse = pseudaminic acid. (iv) Surface electric charge change by E227R. (v) R227 interaction with a neighboring monomer. C) TlpD, chemotaxis receptor for HOCl. HP0559 modeled on PDB 3T9O, the regulatory CZB domain of DgcZ (*E. coli*).

The flagellar filament made of flagellin FlaA (HP0601) (**Figure 6B**, is involved in motility, cell adherence and immune modulation. We have modelled it using the similar *Campylobacter jejuni* homolog ^38^. The flagellin has rod-shaped domains forming a hydrophobic core, and the other domains decorating the surface of the filament are hypervariable. The flagellin is glycosylated by pseudaminic acid at several residues to stabilize the flagellum and to mimic host cell surface, a way for the bacterium to modulate the immune response ^39^, but Fujian-differentiated T364R eliminates one of these sites (**Figure 6B (iii)**). E227R drastically changes the surface electric charge and likely affects its interaction with a neighboring monomer (**Figure 6B (iv)(v)**). Another Fujian-differentiated residue 426 in the conservative core is next to residue 427, which is involved in evasion from TLR5-mediated innate immunity through subunit interaction ^38^. Furthermore, residue 430 adjacent to 426 in the 3-D structure is somehow involved in switching between R and L conformations for swimming/tumbling in chemotaxis in *Campylobacter* ^40^.

TlpD (HP0599, MWE_0916) (**Figure 6C**), is a cytosolic chemotaxis sensor required for colonization. TlpD senses HOCl, an antimicrobial produced by neutrophils during inflammation ^41^. HOCl oxidizes a conserved cysteine (C340) within a 3His/1Cys Zn-binding motif to inactivate chemo-transduction signalling. YunnanMP-differentiated A375S is right by this motif. Additional proteins in Table 1 are described in Supplementary Text and Supplementary figures.

**Table 1.** Genes with a SNP with Fst > 0.5 in the top 20 lists.

| Criterion | Gene | Locus tag | Figure | Category | Annotation | Residue function |
| --- | --- | --- | --- | --- | --- | --- |
| **Fst > 0.6 + top 20 in different pairwise comparisons** | *hefC* | HP0607 | 3B | Efflux pump | Inner membrane component of the multidrug HefABC efflux pump | Channel entrance |
|  | *frpB-4* | HP1512 | 4A | TonB-dependent importer | Outer membrane nickel importer | Ligand binding |
|  | *flaA* | HP0601 | 6B | Motility | Flagellin of flagella, involved in immune system evasion | Glycosylation for host mimicry and more |
|  | *katA* | HP0875 | 6A | Sensor | Catalase sensing and destroying H<sub>2</sub>O<sub>2</sub> | Channel entrance; Dimer-dimer interface |
|  | *hopI* | HP1156 |  | Outer membrane protein | Outer membrane protein HopI |  |
|  | *hopB/alpB* | HP0913 |  | Outer membrane protein | Hop family adhesin HopB/AlpB/Omp21. |  |
|  | *tlpD* | HP0599 | 6C | Sensor | Chemotaxis sensor sensing and destroying HOCl | Active site |
|  | *exbB-2* | HP1339 | S3C | TonB-dependent importer | Energizer/motor in the inner membrane driven by proton | Channel entrance |
| **Fst > 0.6 2 or more SNPs** | *hofC* | HP0486 |  | Outer membrane protein | Hof family protein implicated in adhesion and antibiotics diffusion |  |
|  | *porB* | HP1111 |  | Energy metabolism | Subunit of pyruvate:ferredoxin oxidoreductase, part of the microaerophilic metabolic pathway leading to acetyl~CoA | Subunit interaction |
|  | *gltA* | HP0026 | 5B | Energy metabolism | Citrate synthase, the first enzyme in TCA cycle incorporating acetyl~CoA | Subunit interaction |
|  | *oppD* | HP0250 | 5A | Importer | Oligopeptide ABC transporter, ATPase subunit | Cofactor binding |
| **Fst > 0.5** | *omp18* | HP1125 |  | Outer membrane protein | OmpA family peptidoglycan-associated lipoprotein | Ligand binding |
|  | *fixP* | HP0147 | S4C | Energy metabolism | Subunit of cytochrome c oxidase in aerobic respiration | Channel entrance |
|  | *hypC* | HP0899 |  |  | Hydrogenase expression/formation protein | None (syn) |
|  | *hefD* | HP0971 | 3C | Efflux pump | Outer membrane component of the HefABC multidrug efflux pump | Protein binding |
|  | *copA* | HP1503 | S2E | Exporter | Copper(I) exporter | Protein stability |
|  | *mscS-1* | HP0284 |  | Sensor | Mechano-sensor sensing tension in the membrane | Scaffolding in the periplasm |
|  | *metQ* | HP1564 | S2D | Importer | D-methionine ABC transporter, methionine-binding subunit | Ligand binding |
|  | *frpB-1* | HP0876 | S3A | TonB-dependent importer | Outer membrane heme importer | Ligand binding |
|  | *fecA-1* | HP0686 | S3B | TonB-dependent importer | Outer membrane iron(III) dicitrate importer | Ligand binding; Channel (plug) |
|  | *exbD-2* | HP1340 | S3C | TonB-dependent importer | Inner membrane energizer/motor driven by proton | Channel entrance |
|  | *hpaA* | HP0797 | S3F | Outer membrane protein | Neuraminylactose-binding hemagglutinin | Subunit interaction |
|  | *omp* | HP0358 |  | Outer membrane protein | Putative outer membrane protein |  |
|  | *panD* | HP0034 | S4A (i) | Micronutrient synthesis | Vitamin B5 synthesis | Interaction between cleaved peptides |
|  | *bioD* | HP0029 |  | Micronutrient synthesis | Vitamin B7 synthesis |  |
|  | ***tgt*** | HP0281 | S4A (iii) | Micronutrient synthesis | Q-base synthesis; Q-base on tRNA affects translation accuracy | Active site |
|  | ***rhoD*** | HP1223 | S4B | Detox | Rhodanese detoxifying cyanide generated in microbiome |  |
|  | ***rkiP*** | HP0218 | S2B | Oncoprotein? | Mimic of human RKIP tumor suppressor | Interaction with<br>signal peptide |
|  | ***hcpX*** | - | S2A | Effector | SLR family with repeated alpha-helix pairs | Human protein<br>binding |
|  | ***dsbI*** | HP0595 | S3D | Secretion | S-S formation | Active site |
|  | ***jag*** | HP1451 |  | Secretion | Regulator of VirB11/Cag-alpha gate of Cag secretion system | Cofactor binding |
|  | ***lpxE*** | HP0021 | S2C | Membrane lipid modifier | Lipid A 1-phosphatase to hide it from the innate immune response | Substrate binding;<br>Active site |
|  | ***cfaS*** | HP0416 | S3E | Membrane lipid modifier | Cyclopropane-fatty-acyl-phospholipid synthase for acid protection | Substrate binding |
|  | ***fur*** | HP1027 | S4D |  | Regulator of Fe/Ni import, redox balance and acid response | Cofactor binding |

## Discussion

Analyses of *H. pylori* in East Asia have tended to emphasize their homogeneity and uniformly high virulence potential. A single subpopulation of the bacteria, hspEAsia, is prevalent in the region. The hspEAsia strains have been found to be invariably CagPAI positive, with the *cagA* gene containing the ABD EPIYA motif which is thought to promote strong binding of the protein to SHP-2 ^42^. HspEAsia strains also characteristically have a particular variant of *vacA*. Our large collection of genomes of *H. pylori* from multiple regions in China, Japan and Korea confirm these observations. Only a small number of these isolates, of which a majority of are from Mongolia and Okinawa, belonged to other *H. pylori* populations. For the scope of this study, these were excluded from subsequent analyses. All but 13 of the total 381 isolates were *cagA* positive and 355 have the characteristic ABD EPIYA type, including 316 out of 329 hpEAsia isolates.

Our results add a layer of complexity to the picture of uniformity by demonstrating that there is differentiation of *H. pylori* strains of the hspEAsia subpopulation between regions in East Asia. Despite a large burden of gastric disease in the region, most *H. pylori* infections by hspEAsia are asymptomatic, and the gastric cancer incidence varies widely across the region, especially in China. A large part of these differences might be attributable to diet and environment. However, our results imply that there are also bacterial factors differentiating these regions, which may be significant for disease development, especially because the bacteria themselves can adapt to environmental conditions ^5^.

Geographic differentiation between populations accumulates progressively when migration rates between them are low. Further, adaptation of bacteria to differences in environmental conditions can greatly accelerate the process of differentiation in specific regions of the genome. Our results, in combination with a previous study of genetic variation within the Americas ^29^ suggest that there are a handful of loci that have undergone rapid differentiation in several regions, and therfore may be considered keys for host adaptation. These include the genes *frpB-4, hefC, alpB/hopB*, and *hofC*.

Our strategy to identify geographically differentiated SNPs by dividing one population (hspEAsia) into minimal subpopulations, i.e. strains with consistent population and strain labels, and comparing fixation index (Fst) site-by-site between these populations reveal numerous loci of differentiation (**Suppl. table S4**). Of these we discuss the 12 with the strongest evidence for being involved in local adaptation in more detail in the main text in this paper. Some of the region-specific SNPs are in genes encoding for proteins that have been implicated host interaction and virulence in the narrow sense: attack by immune system (catalase, TlpD), host adhesion (HopB/AlpB and several outer membrane proteins), and host surface mimicry (the flagellin FlaA). Several of the other genes are transporters that may have implications for antimicrobial resistance, or are involved in nutrient acquisition. These results suggest that various host-adaptive changes in many host-interaction proteins lead to population differentiation. A similar gene set was found when rapid genome changes were investigated in shorter-term, intra-body micro-evolution ^43^.

Epidemiological and experimental evidence suggests that iron-deficiency increases *H. pylori* virulence and risk of gastric cancer ^44^ In our analyses we could see both in iron and nickel metabolism highlighted by regional changes. Apart from the above-mentioned *frpB-4*, genes encoding for the TonB motor proteins ExbB-2 and ExbD-2, heme transporter *frpB-1 ^45^* appear in the list of the 34 genes with Fst >0.5 and the central transcription factor ferric uptake regulator, *fur* had regional variants (Supplementary text). The causal mechanism underlying this association is not clear but it plausibly reflects bacterial response to nutrient limitation. Simply put, it is possible that bacteria adopt more aggressive strategies in interacting with the host and its microbiome when iron and other metals such as nickel, which is necessary for urease function, is in short supply.

In other organisms, linkage means that high Fst regions often occur in large blocks, making it difficult to infer which sites are involved in local adaptation. However, *H. pylori* lineages recombine with each other, exchanging substantial fraction of their DNA in individual mixed infections ^46^. The size of replacement in one event can be as short as 40 bp^47^, with the result that linkage is broken down rapidly. This means that individual nucleotides can rise to high frequencies in specific populations, meaning that local adaptation can potentially occur on a very exquisite scale.

Many of the highly differentiated amino-acid changes are close to critical residues of the protein and are plausible candidates to cause important functional changes, based on 3-D modelling and previous functional analyses. We have suggested possible functional consequences but validation by targeted experiments and clinical observations is necessary. Although the functional consequences of genomic differentiation of *H. pylori* within different parts of the world remain to be elucidated, the presence of this differentiation already has potential clinical utility. All else being equal, individuals who are infected by *H. pylori* that are characteristically found in high gastric cancer incidence regions are likely to be at higher risk than those associated with lower incidence regions, firstly because the bacteria may be more virulent but secondly because infection to the bacteria might also be a marker for exposure to environmental factors that underlie the high disease risk.

## Acknowledgements

We thank Jonas Korlach and Primo Baybayan for help in SMRT sequencing. We would like to thank and the staff of Comparative Genomics Laboratory at National Institute for Genetics, Mishima, Japan for supporting genome sequencing. We thank Dr Chao Yang, Professor Yujun Cui and Ruifu Yang for the discussions about *H. pylori* population genetics of *H. pylori*, and M. Zwama for discussion on HefC, Mizuki Ohno and Yosuke Kawai for discussion on human genetics. This work was supported by a grant from the State Key Laboratory of Infectious Disease Prevention and Control (SKLID) (2014SKLID102) of the Chinese Center for Disease Control and Prevention. Supported by National Science and Technology Major Project (2018ZX10712-001) and a joint project ‘Isolation and sequence analysis of *Helicobacter pylori* strains collected from investigations on carriage rate ‘. KT was supported by Swedish Society for Medical research (SSMF). Parts of the bioinformatic analyses were performed on resources provided by Swedish National Infrastructure for Computing (SNIC) through Uppsala Multidisciplinary Center for Advanced Computational Science (UPPMAX) under projects snic2018-8-24/uppstore2017270, partially funded by the Swedish Research Council through grant agreement no. 2018-05973. This work was also supported in part by MEXT KAKENHI (19K22543, 17H04666, 26113704, 25291080, 221S0002 to IK, 18K14766 to YK) and by Shanghai Municipal Science and Technology Major Project (No. 2019SHZDZX02 to DF).

Note to the cover page

TEAMHp (Team for East Asian Genomics of *Helicobacter pylori*) represents the following people.

Takahiro Bino

NIBB Core Research Facilities, National Institute for Basic Biology, Okazaki, Aichi, Japan Data Analysis

Masaki Fukuyo

Department of Molecular Oncology, Graduate School of Medicine, Chiba University, Chiba 260-8670, Japan, Kazusa DNA Research Institute Chiba 292-0818, Japan

Rumiko Suzuki

Department of Environmental and Preventive Medicine, Oita University Faculty of Medicine Provide strains

John Harting

Pacific Biosciences, Menlo Park, CA 94025, USA Data Analysis

Mototsugu Kato1,2

1 Division of Endoscopy, Hokkaido University Hospital, Sapporo, Hokkaido 060-8468, Japan.

2 Department of Gastroenterology, National Hospital Organization Hakodate Hospital, Hakodate, Hokkaido, Japan.

Provided strains

Mutsuko Konno

Department of Pediatrics, Sapporo Kosei General Hospital, Sapporo, Hokkaido, Japan Provided strains

Yuji Kohara

Advanced Genomics Center, National Institute of Genetics, Shizuoka 411-8540, Japan Carried out Pacbio sequencing

Christine Lambert

Pacific Biosciences, Menlo Park, CA 94025, USA Carried out experiments

Yohei Minakuchi

Comparative Genomics Laboratory, National Institute of Genetics, Shizuoka 411-8540, Japan

Carried out experiments.

Shin Nishiumi

Department of Gastroenterology, Graduate School of Medicine, Kobe University, Chuou-ku, Kobe, Hyogo, 650-0017, Japan.

Provided strains

Shuji Shigenobu

(NIBB Core Research Facilities, National Institute for Basic Biology, Okazaki, Aichi, Japan) Data Analysis

Noriko Takahashi

Department of Computational Biology and Medical Sciences (formerly Department of Medical Genome Sciences

Graduate School of Frontier Sciences, University of Tokyo, Tokyo 108-8639, Japan, Institute of Medical Science, University of Tokyo, Minato-ku, Tokyo 108-8639, Japan,

Kyorin University School of Medicine, Mitaka, Tokyo 181-8611, Japan,

Atsushi Toyoda

Comparative Genomics Laboratory and Advanced Genomics Center, National Institute of Genetics, Shizuoka 411-8540, Japan

Carried out Pacbio sequencing

Ikuo Uchiyama

Laboratory of Genome Informatics, National Institute for Basic Biology, National Institutes of Natural Sciences, Okazaki, Aichi, Japan

Data Analysis

Hirokazu Yano

Department of Computational Biology and Medical Sciences (formerly Department of Medical Genome Sciences),

Graduate School of Frontier Sciences, University of Tokyo, Tokyo 108-8639, Japan, Institute of Medical Science, University of Tokyo, Minato-ku, Tokyo 108-8639, Japan,

Graduate School of Life Sciences, Tohoku University, Sendai 980-8577, Japan

Carried out experiments

Masaru Yoshida

Department of Gastroenterology, Graduate School of Medicine, Kobe University, Chuou-ku, Kobe, Hyogo, 650-0017, Japan.

Provided strains

## Conflict of interest

Christine Lambert and John Harting are full-time employees at Pacific Biosciences of California, a company developing single-molecule sequencing technologies. The other authors declare that they have no conflict of interest.

## Notes

https://www.ncbi.nlm.nih.gov/bioproject/PRJNA482300

### Competing Interest Statement

The authors have declared no competing interest.

