## Supplementary figures and text for "Genomic differentiation within East Asian *Helicobacter pylori*"

### Supplementary materials

#### Supplementary Table S1. Geographical coordinates of isolate locations
#### Supplementary Table S2. Isolate details
#### Supplementary Table S3. Reference isolates
#### Supplementary Table S4. Details of subpopulation-specific SNPs.

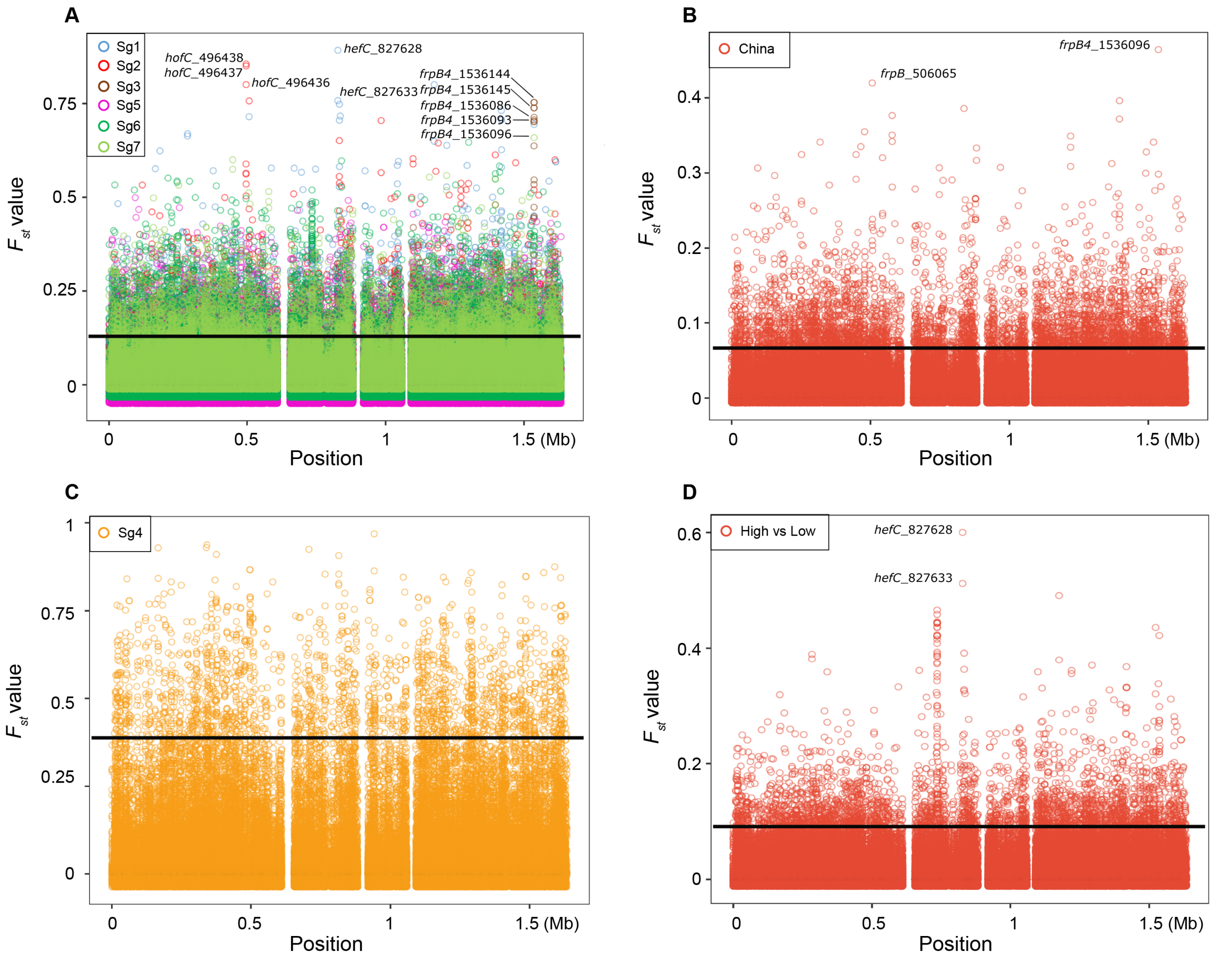


***Supplementary Figure 1.*** Statistics of Fst value for the subgroups.

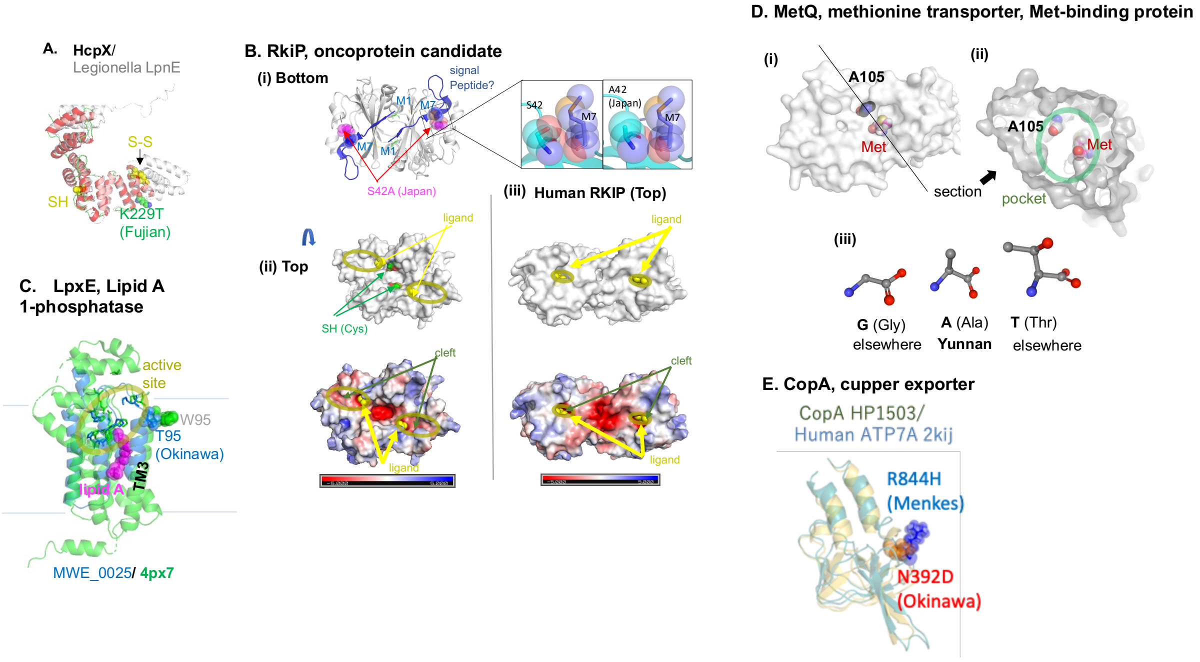


**Supplementary Figure 2**. A) HcpX. MWE_1300 modeled on LpnE (PDB 6DEH). B) RkiP, an oncoprotein candidate. (i) RkiP (= HP0218) structure (PDB 2evv) before signal-peptide excision and S-S bonding showing interaction of S42 with M7 of the hypothetical signal peptide and its alteration by Japan-specific A42 (modeled by PyMol). (ii) The bottom figure shows electric charge distribution on the surface (PyMOL). (iii) Human RKIP (PDB 1beh) with S-S bond with a bound ligand (cacodylate). C) LpxE, lipid A 1- phosphatase (HP0021). MWE_0025 modelled on PDB 4px7 (PgpB of *E. coli*). D) MetQ. (i, ii) MWE_1564 modelled on and aligned with PDB 4yah (*E. coli* homolog). (iii) Diverse amino acids at residue 105. E) CopA. MWE_1691, a well-conserved copper(I) exporter of the P-type ATPases characterized by a P~protein intermediate. Okinawa-specific N392D is in the actuator domain involved in structural change between the open and closed configuration. A mutation in the corresponding site in its human homolog ATP7A causes protein disappearance and leads to Menkes syndrome affecting copper metabolism 7.

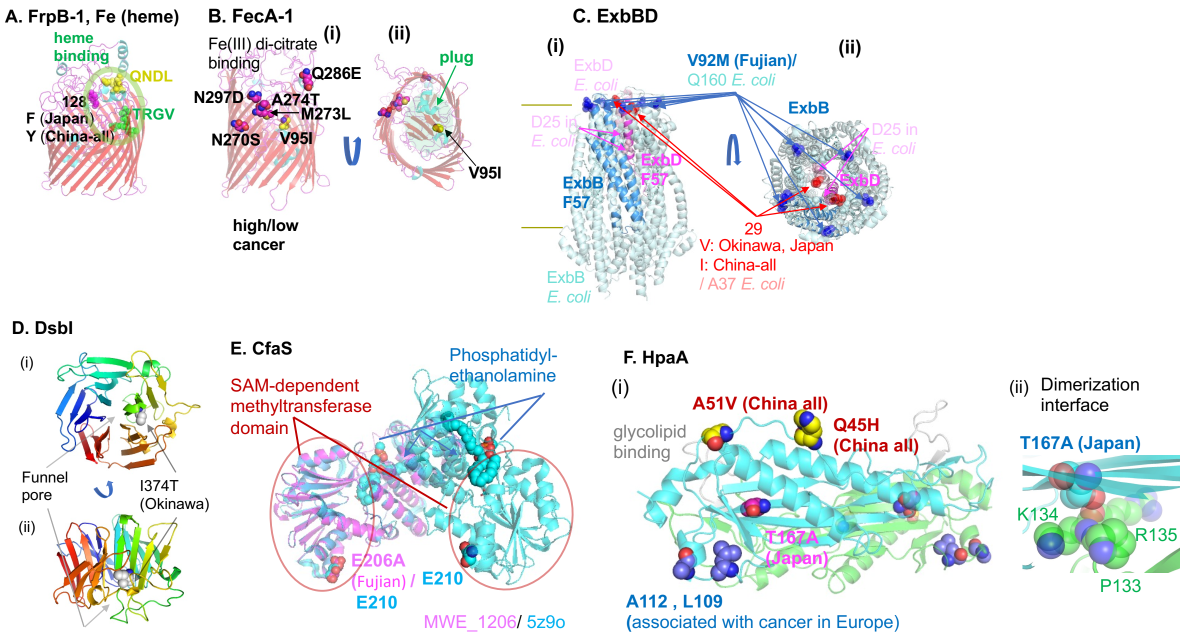


**Supplementary Figure 3**. A) FrpB-1. MWE_0564 modelled on PDB 4aip. 26695 carries *frpB-1* (HP0876), *frpB-2/3* (HP0916/5) and *frpB-4* (HP1512). B) FecA-1. MWE_0823 (an HP0686 homolog) modelled on PDB 1kmp (FecA of *E. coli*). The residues distinguishing the low stomach cancer populations and the others are in spheres. C) ExbBD motor. HPF57_1294 and HPF57_1295 were modeled on PDB 6tyi (*E. coli*) with ExbB:ExbD = 5:2 stoichiometry. (MWE_1546 is a pseudogene.) One F57 ExbB molecule is in blue is on five *E. coli* ExbB subunits in pale blue. One F57 ExbD molecule is in magenta is on two *E. coli* ExbD molecules in light pink. Residue 92 in F57 ExbB (113 in HP1339) and corresponding Q160 residues in E. *coli* ExbB are in deep blue. Residue 29 in HPF57_1295 and corresponding A36 residues in *E. coli* ExbD are in red. D25 in the proton channel of *E. coli* ExbD is in spheres. D. DsbI. C-half (166-) of MWE_0921 was modeled on PDB 3qqz (YjiK in *E. coli*) as a 5-blade beta-propeller. E. CfaS. Cyclopropane fatty acid synthase. MWE_1206 modeled on PDB 5z9o (*Lactobacillus acidophilus* homolog). F. HpaA. MWE_0650 modelled on its paralog HP0492 (PDB 2i9i), not its ortholog HP0797. The association is from a GWAS.

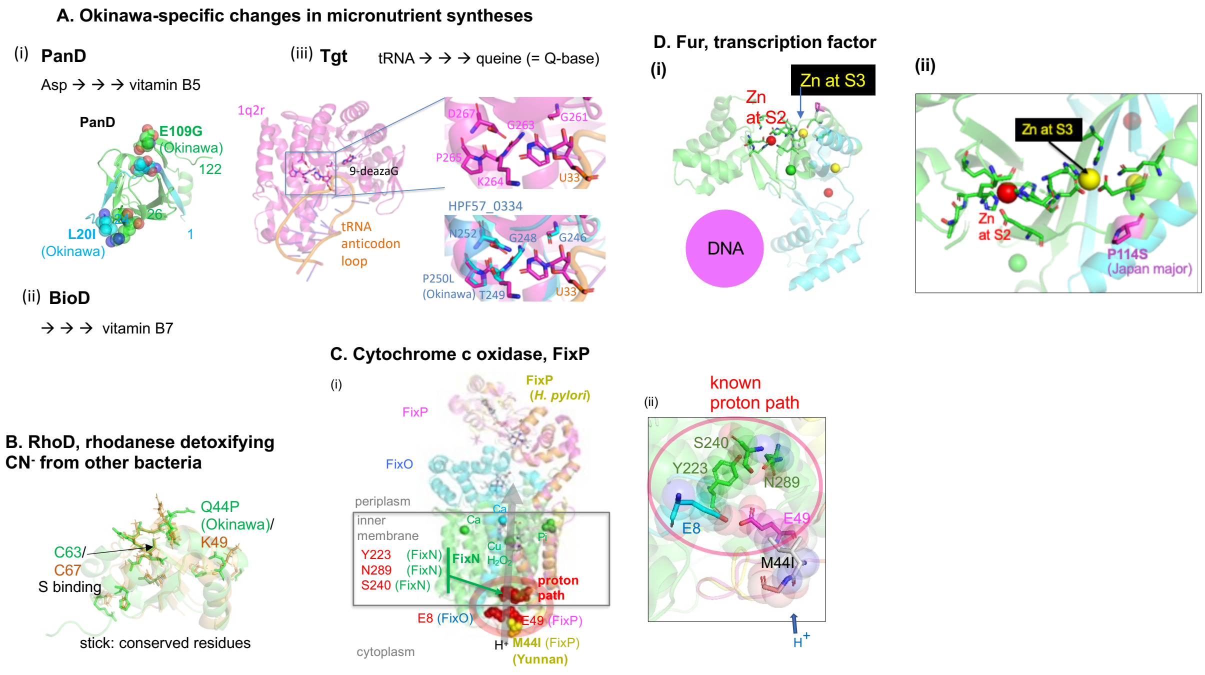


**Supplementary Figure 4.** A) (i) PanD, Aspartate 1-decarboxylase. Solved structure of HP0034 (PDB 1uhe). A monomer unit out of the tetrameric enzyme is shown. (ii) BioD. Dethiobiotin synthetase. (iii) Tgt. Queine-tRNA ribosyltransferase. HPF57_0334 was modeled on PDB 6gym (*Zymomonas mobilis* homolog with W95F mutation) and aligned with PDB 1q2r (the same protein without a mutation).. B) RhoD. MWE_1424 modeled on HP1223 (2k0z)/PspE (PDB 2jtq). C) FixP in cytochrome c oxidase. MWE_0216 (HP0147 homolog from *E. coli*) modeled on, and aligned with, Pseudomonas FixNOP (PDB 3mk7). The five known proton channel component residues are in spheres. D). Fur. PDB 2xig.

### Supplementary text. Function and role of population-differentiated amino-acids not discussed in main text.

#### Sensors and transporters

##### *Mechanosensor.*

HPF57_0337, **MscS-1,** includes a **MscS** (**mechanosensitive channel of small conductance)** family member [1]. When a hypo-osmotic shock raises the pressure in the cell, MscS family senses the increase of tension in the membrane by the force from the lipids and releases solutes and water and plays various roles in signal transduction. Fujian-specific H139R is predicted to be on coiled-coil helix, often seen in a periplasmic scaffold, N-distal to the MscS domain. This MscS family member might have a novel type of host interaction.

##### *Methionine importer*

Amino acids and oligopeptides are important nutrients for *H. pylori* and other members of the microbiome as well as for the host human body. MetQ (HP1564, MWE_1780), methionine-binding subunit of a D-methionine ABC transporter (**Fig. S2D**), has Yunnan-specific TG105A in the Met binding pocket. Oxidative damage of Met on proteins by host-driven oxidative stress is a serious problem for *H. pylori* [2]. The dependence of cancer cells on exogenous methionine is known as the Hoffman effect [3].

##### *TonB-dependent importers*

*H. pylori* reference strain 26695 carries the following *frpB* homologs: frpB-1 (HP0876), frpB-2/3 (HP0916/5), and frpB-4 (HP1512). **FrpB-1** for heme import carries a site (128) distinguishing between Japan (F) and China-all (Y) near its two heme binding motifs, QNDL and TRGV [4] (**Fig. S3A**).

**FecA-1** imports iron(III) dicitrate (**Fig. S3B**). It determines the survival of *Helicobacter pylori* in the stomach [5]. Comparison of high cancer areas vs. low cancer areas revealed four clustered amino-acid changes on its upper side and one in the plug with Fst values of 0.44-0.48.

**TonB-motor components ExbB-2** and **ExbD-2** [6] carry Fujian-specific V92M and residue 29 distinguishing Okinawa/Japan (V) and China-all (I), respectively, at the periplasm-inner membrane interface (**Fig. S3C**).

##### *Exporters*

**CopA** (MWE_1691), a well-conserved copper(I) exporter of the P-type ATPases characterized by a P~protein intermediate, has Okinawa-specific N392D in the actuator domain involved in structural change between the open and closed configuration. A mutation in the corresponding site in its human homolog ATP7A causes protein disappearance and leads to Menkes syndrome affecting copper metabolism (**Fig. S2E**) [7].

#### Outer membrane proteins

**HpaA** (**Fig. S3F**) is an adhesin well studied as a virulence factor. It has Japan-specific T167A and China-specific Q45H and A51V as well as gastric-cancer associated L109 and A112 (detected in GWAS) [8] in the glycolipid-binding ends [9]. T167A is also at the interface between two identical subunits (**Fig. S3F (ii)**).

**OmpA** family are outer membrane proteins non-covalently anchored to peptidoglycan and can form 8-stranded beta-barrel in the N-terminus. They have pathogenic roles including adhesion, invasion, and intracellular survival as well as evasion of host defenses and stimulation of pro-inflammatory cytokine production [10]. **Omp18** (HP1125) lipoprotein likely of this family is involved in persistent colonization by evading interferon γ signaling [11]. **HP0358** with Japan-specific 151 is another protein likely of OmpA family.

#### Host interaction factors

##### *RkiP, an oncoprotein candidate mimicking human RKIP*

This protein (HP0218) (**Fig. S2B**) belongs to PEBP family functioning in lipid binding and regulation of signaling pathways. Its human member RKIP (**Fig. S2B (iii)**) is a tumor suppressor inhibiting MAPK pathway of signal transduction leading to stomach cancer.. In HP0218's solved structure, likely representing a cytoplasmic precursor form, the two monomers have strong interaction only around C65 (and N-terminus, see the paragraph end) from each monomer in the parallel configuration just as the human homolog RKIP. All the other structurally characterized bacterial members have anti-parallel configuration instead. Bonding of the SH pair into S-S would lead to a shape and an electric charge distribution around the ligand binding area closely resembling the human RKIP's. We hypothesized that *H. pylori* member has evolved a mimic of human RKIP and named them RkiP (from *rkiP* gene). *H. pylori* infection increases in RKIP phosphorylation [12] and RKIP enhances cell death in infection [13]. Expression of RkiP is altered by adhesion to human cells [14]. We hypothesize that RkiP interferes with RKIP action as a mimic, which results in increase in RKIP phosphorylation and switches MAPK pathway to oncogenesis. RkiP carries a weak signal of signal peptide (SignalP-5.0) in a region corresponding to the signal peptide of YbcL of *E. coli*. Japan-specific S42A, modeled by *in silico* mutagenesis (PyMOL), alters interaction of residue 42 with M7 on this putative signal peptide (**Fig. S2B (i) right**). We hypothesize that this change may affect maturation and the above function of RkiP.

##### *HcpX, an effector candidate*

**HcpX** (**Fig. S2A**) belongs to SLR protein family, a subfamily of tetratricopeptide repeat (TPR)-containing proteins. They contain a motif of two antiparallel alpha-helices such that its tandem arrays generate a right-handed helical structure with an amphipathic channel that accommodate the complementary region of a target protein. HcpC of this family (HP1098) is secreted and interacts with several human proteins [15]. HcpX, carrying only one S-S bond connecting the helix pair, is well modeled on LpnE, a Legionella effector targeting human inositol polyphosphate 5-phosphatase, OCRL-1 [16]. Fujian-specific K229T changes surface shape and surface electric charge presumably affecting its interaction with target proteins.

##### *Jag regulating Cag secretion system*

VirB11 (= Cag-alpha = HP0525) ATPase of Cag secretion system forms a hexamer-ring gate to transport CagA and other molecules to human cells [17]. **Jag** is its unlinked (mapping outside of CagPAI) negative regulator forming its partial lid [18]. Fujian-specific S207N is in its S8-like nucleic-acid binding domain.

##### *DsbI*

Dsb (disulfide-bond formation) proteins receive electrons generated by S-S bond formation in the cytoplasmic membrane. **DsbI** [19] (HP0595, MWE_0921) forms a subfamily of DsbB in epsilon-proteobacteria. *H. pylori* DsbI is required for colonization in mice [29]. It has Okinawa-specific I374T by the funnel of its predicted 5-balade beta-propeller (**Fig. S3D**) in its C-half.

##### *Modifiers of membrane lipids*

Kdo2-lipid A is an essential component of lipopolysaccharide but stimulates the innate immune response. **LpxE, lipid A 1- phosphatase** (HP0021, MWE_0025) (**Fig. S2C**) removes its 1-phosphate to camouflage it and also to confer resistance to host-derived cationic antimicrobial peptides (CAMPs) and to antibiotics polymyxin [20]. Okinawa-specific T95A is by the active site and at the top of a helix that catches the lipid.

The lipids containing cyclopropane-fatty-acid protect bacteria from acidity and antibiotics [21]. **CfaS, cyclopropane-fatty-acyl-phospholipid synthase (**HP0416), transfers a methylene group from S-adenosylmethionine across the *cis* double-bonds of unsaturated fatty acyl chains for their synthesis. It has Fujian-specific E206A at the end of its methyltransferase domain (**Fig. S3E**).

#### Metabolic enzymes

##### *Vitamin B5 and B7 syntheses in Okinawa*

Enzymes for synthesis of three micronutrients are altered in *H. pylori* of Okinawa subgroup (**Fig. S4A**). **PanD** for pantothenate (**vitamin B5**) synthesis (**Fig. S4A (i)**) is self-processed to produce two chains, which stay together by interaction at Okinawa-specific L20I and E109G. **BioD** for biotin (**vitamin B7**) synthesis carries Okinawa-specific N97D.

A GWAS of host species specificity of *Campylobacter jejuni* revealed that vitamin B5 synthesis genes were conserved in cattle isolates than in chicken isolates likely because this vitamin is present in the grass diet of the cattle [22]. Likewise, alteration of the enzymes for synthesis of the three micronutrients (for Q-base, see below) in Okinawa might be related to the unique human diet in Okinawa islands [23].

##### *Q-base synthesis in Okinawa*

**Tgt** (HPF57_0334) (**Fig. S4A (iii)**), queuine tRNA-ribosyltransferase, replaces G34 in the anticodon of several tRNAs by a precursor of quine (Q-base), which modifies codon recognition and affects efficiency and accuracy of translation [24]. Hosts and members of their associated microbiota compete for the salvage of Q-base precursor micronutrients [25]. Tgt contributes to virulence of *Shigella flexneri* and *E. coli* [26]. Okinawa-specific P250L is by the residues recognizing U33. This residue is tightly attaching to of the three active site residues.

The differentiation in synthesis of Q-base may affect carcinogenesis. Decreased Q-modification of tRNA has been demonstrated for a large number of neoplastic tissues. Q-base modification of tRNA alters translational fidelity. The proteome of cancer cells is not fully encoded by their transcriptome, although the contribution of (mis)translation to such diversity remains to be elucidated [27]. The possibility that Q-base-mediated interaction between *H. pylori* and human cells promotes carcinogenesis and explains low cancer incidence in Okinawa is testable at several levels.

#### Fur transiption factor

Fe and Ni import and homeostasis as well as redox homeostasis and acid response are regulated by **Fur** (Ferric uptake regulator), a transcription factor required for colonization (**Fig. S4D**). Its Japan-specific P114S is close to Zn at S3 important in DNA binding [28].
